# ProLego: tool for extracting and visualizing topological modules in protein structures

**DOI:** 10.1101/225565

**Authors:** Taushif Khan, Shailesh Kumar Panday, Indira Ghosh

## Abstract

**Background:** In protein design, correct use of topology is among the initial and most critical feature. Meticulous selection of backbone topology aids in drastically reducing the structure search space. With ProLego, we present a server application to explore the component aspect of protein structures and provide an intuitive and efficient way to scan the protein topology space.

**Result:** We have implemented in-house developed “topological representation” in an automated-pipeline to extract protein topology from given protein structure. Using the topology string, ProLego, compares topology against a non-redundant extensive topology database (ProLegoDB) as well as extracts constituent topological modules. The platform offers interactive topology visualization graphs.

**Conclusion:** ProLego, provides an alternative but comprehensive way to scan and visualize protein topology along with an extensive database of protein topology.

ProLego can be found at http://www.proteinlego.com

## Background

Understanding of protein fold universe remains one of the major goal in post genomic era. Numerous attempts in exploring the nature of protein structure space led to classification schemas like SCOP [1], CATH [2], ECOD [3] PCBOST [4] that investigate protein’s structural, functional and evolutionary features. Topology based approach has been recently exploited to examine the structure space of proteins and provide insights into fold designing and evolution [4–7]. With the increasing sophistication in experiments, number of topology can be tested in high-throughput screening to analyse the global determinant of folding and stability[8]. Moreover, topology has been considered to be exceptionally convenient to address the nature of folding profile by both experimental and computational approaches [9]

Structural modularity is crucial in conferring functional and structural diversity of proteins [6, 10]. This concept can be explained using analogy of structural modules to “Lego” blocks that can be reused to build proteins with tailored functionality. Although the analogy with “Lego” blocks might oversimplify the complex nature but can depict well the current knowledge space [11]. Protein topology has been studied using several graph-based techniques to understand domain arrangement [12, 13], protein folding pathways [9], analysis of different biochemical activities and structural comparison[14]. With the emergence of computer graphics, protein topology representation evolved from manual drawing [15] to scalable graphics representation [16, 17]. However, only handful of methods are available that provide automatic generation of the protein topology diagram (Table S1). The most recent addition in the list is PTGL [17], which in addition to different protein folding graphs also able to address the ligand interaction with protein topology. However, the issue of module identification and visualisation could be addressed in much efficient way as proposed by protein lego server, as reported here.

With ProLego, we propose a platform that can be used to analyse protein topology and its modular architecture. ProLego, along with generating improved topology cartoon diagrams, provide tools for searching proteins with similar topology and extracting constituent structural modules. With the implementation of protein “topology string” on non-redundant protein chains, we propose a protein topology database (ProLegoDB) focusing on the composition and organisation of secondary structures.

## Implementation

ProLego is a “pythonic” solution to the topology generation with the help of D3.j s (a JavaScript library) for visualization. Briefly, in the background, user provided protein chain examined for secondary structure (SS) contacts and relative orientation. The SS-contact definition is considered based on the presence of corresponding residual contacts (as in [14, 18]). From proteins atomic coordinate, an adjacency matrix of SS-contact has been generated, from which 1D “topology string” has been built. The “topology string” encompasses the composition, contact and relative arrangement of SS (see Supplementary section 1.2).

The present study implements a pipeline that uses above mentioned (a) “Topology String” (or Contact String) to define relative position and orientation of secondary structure elements (SSE; DSSP definition [19]), (b) a database “ProLegoDB” of pre-calculated topology information of representative proteins [10] and (c) provides a topology visualization platform. “Topology String” translates protein topology in an intuitive character string, which makes searching and storing topologies efficient. The architecture of the server is discussed in supplementary (Figure S2 and Section S1.3)

## Results

ProLego leverages the component approach of protein topology space to extract inherent modules, similar topology and assigns topology frequency class (Preferred, Non-preferred). Representing protein topology as a graph of secondary structure, ProLego provides visualization focusing on different representation (Fig 1A). The ProLegoDB is an extensive database of protein topology generated by analysis of representative datasets. The proposed web platform provides the access to explore the topology space. The backend use of string-based (“topology string”) search method makes the process efficient and intuitive. In the following section, some of the key finding in nature of topology space by analysis of different representative non-redundant datasets have been discussed. Some of the salient features of the server application has been presented along with a comparative study with current state-of-the-arts topology servers.

**Figure 1:**
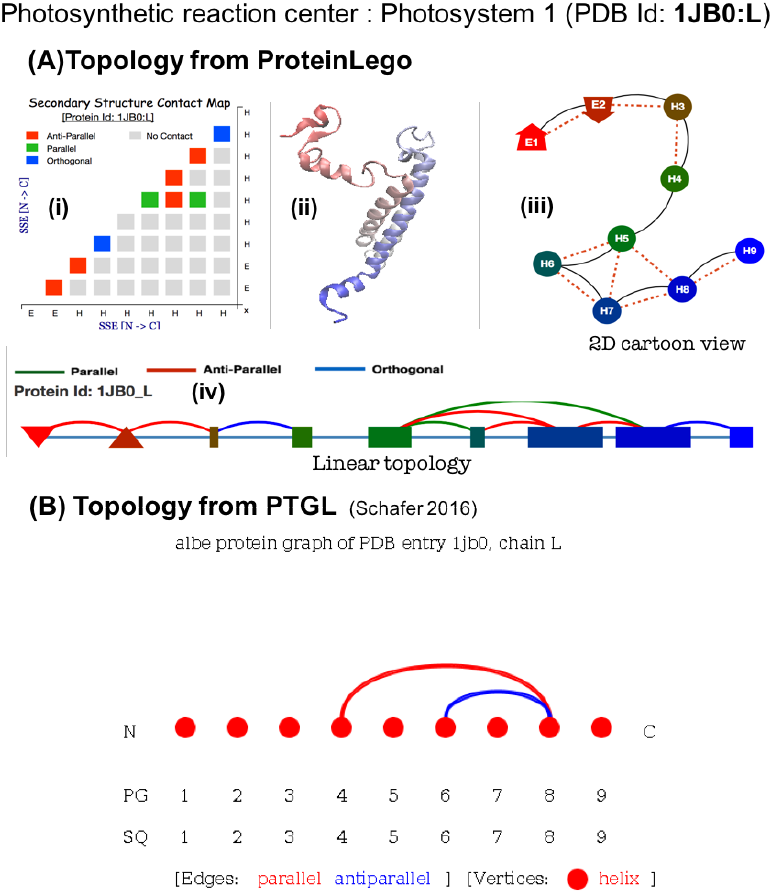
Comparing topology visualization using ProLego (A) and PTGL(B) and for the case of photosynthetic reaction centre (Photosystem1 (PDB Id: 1JB0; chain: L)). The chain has seven alpha-helices and an anti-parallel beta sheet at the N-terminal. Fig A.ii, shows a cartoon view of the protein chain is generated using VMD, colour coded in red to blue representing N and C terminal respectively. In linear topology (A.iv) strands are represented as triangles (with relative orientation as up/down triangle) and helices are represented as rectangle. The length of helical rectangles scaled as per number of residues in the helix. The protein chain is represented as red to green to blue as passes from N to C terminal. The linear lines, connecting secondary structure (SS) blocks shows the chain connectivity, whereas the arc lines represent the spatial connectivity and type of SS contact (colour coded as labelled in Table S4). The secondary structure contact map (A.i), shows all spatial contact between pairs of SS. A 3D carton representation (VMD generated A.ii) and 2D topology cartoon (A.iii) plot is generated from ProLego. The 2D ProLego cartoon shows contact between two SS blocks by red dotted lines and chain connectivity by black continuous line. Figure B, shows the topology representation of the same protein generated using protein topology graph library (<u>http://ptgl.uni-frankfurt.de/results.php?q=1jb0L</u>), the alpha-beta graph. The graph represents SSEs from N to C terminal in left to right fashion. Helices are represented as circles and stands as rectangles. PTGL considers, 3_10_ helices also in total helix, hence the addition of 1st and 7^th^ helix, giving total number of helix to 9 instead of 7 alpha-helix as per ProLego in this protein. PTGL misses the N-terminal sheet, which is represented as up-down triangle (for anti-parallel orientation) in case of ProLego.

### Distribution of proteins in topology space

Using secondary structures (SS) as building blocks of protein structures, we have defined topology as the arrangement, spatial contacts and organisation of SS in a protein chain. Applying this simple but efficient definition, we have scanned representative protein structure databases and extracted underlying topological space. The representative data sets have been curated for sequence redundancy with state-of-the-arts methods to mitigate the effect of structural bias in current protein structure space. We have examined a total of 58186 protein chains from PDB and 14408 protein domains from curated domain databases. As the study undertakes the composition of SS, we have grouped the whole structure space with different composition of SS (or “structure class”) as all-alpha (A), all-Beta (B) and mix Alpha-Beta (AB) referred as. A detail of dataset statistics can be read from Table 1.

**Table 1:** Description of ProLegoDB.

| Structure Class | Topologies<sup>*</sup> | Proteins<sup>a</sup> | Domains<sup>b</sup> |
| --- | --- | --- | --- |
| A | 3315 | 6064 | 2134 |
| B | 2485 | 3955 | 1520 |
| Mix AB | 1401 | 48167 | 10754 |
| Total | 7201 | 58186 | 14408 |

The topology database, ProLegoDB, describes protein topology space. Representative datasets of non-redundant protein chains and domain has been constructed as described in (S1.3). Above table summarises the database with different structure class (A: all-alpha, B: all-beta and mix AB: Alpha-Beta). Number of <sup>\*</sup>statistically significant topology group for each structure classes has been shown with table heading of “Topologies”. Number of proteins in the database for each structure class has been reported in the next columns. <sup>a</sup>Protein chains are considered from extracted non- redundant datasets of PDB, whereas <sup>b</sup>Domains are protein entry from curated domain databases of CATH (3.5) and Astral (SCOP v1.75). The maximum pairwise sequence identity between chains are < 40%.

Distribution of proteins in different topologies have been examined and statistically significant topologies are identified (P-Value < 0.001). The significance are further examined with restricting false discovery rate to less than 0.1%, using P-Value correction method [20] (Table S5). Figure 2, describes the distribution of proteins in statistically significant topologies. We have compared the topology and protein space in “Prevalent” (P) and “Non-prevalent” (NP) classes. For each case, the density of distribution is represented by the width of the violin plot and the spread of the inter-quartile region describes the variation. Comparing the density distribution, a clear distinction in distributions of “P” and “NP” can be observed. For each case, maximum density of the data can be found around their respective mean and interquartile regions, whose values varies for topology and proteins in both cases. Examining the distributions, it can be observed that the topologies in “P” are only ~ 20% of the total topology space, whereas it caters to ~70% of total proteins, which is reverse in the case of “NP”. This characteristic of distribution for topology is quite evident, however proteins have subtle higher variance, distributed around mean of ~60% for “P” and ~40% for “NP”. Similar analysis has been performed for different datasets and among structural classes (Figure S5). Among all studied cases, we have observed the consistent distribution of topology space, with tolerable variance in protein distribution across structure classes.

**Figure 2:**
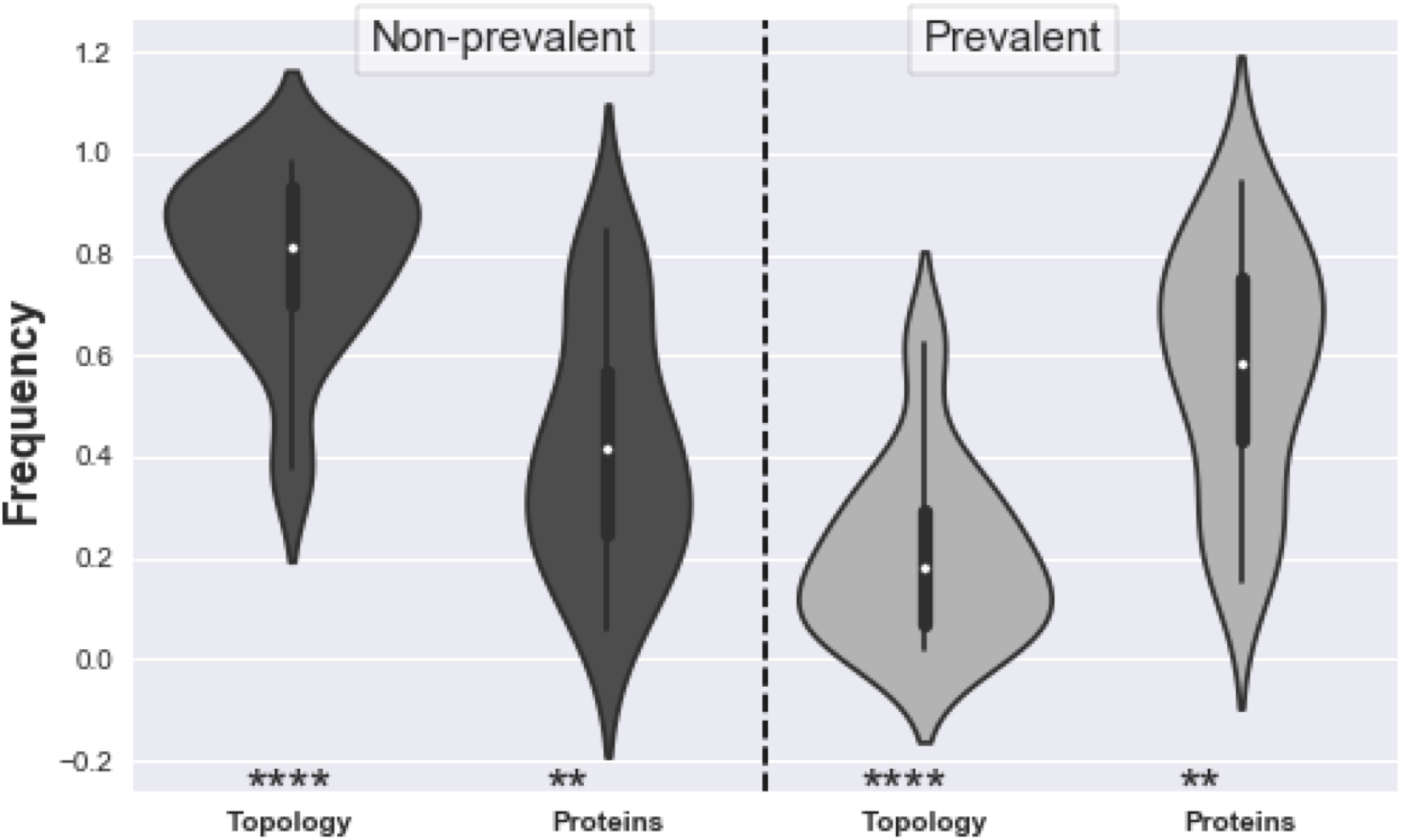
Distribution of topology and protein in groups of “Non-Prevalent” (left to dashed line) and “Prevalent” (right to dashed lines) has been shown as violin plots. This plot is generated for the statistically significant topologies (P-value < 0.001; Table S3), from represented dataset of PDB (58186 protein chains). Description of dataset has been provided in the text and supplementary. The shape of violin plot describes the kernel density estimation of the distribution of data in different topologies and proteins. A summary of statistics can be drawn from the inner boxplot. The white dot represents the median, thick bar shows the interquartile range and thin line describes the 95% confidence interval. A clear distinction can be drawn on the nature of distribution of proteins as well as topologies in “Prevalent” and “Non-prevalent” groups. A comparison of distribution with non-parametric Wilcoxon rank-sum test has been performed and P-values are indicated as ‘*’ (‘****’:P-Val < 0.001 and ‘**’: P-val < 0.01) in the bottom.

Using the topology string, it is possible to draw a distribution and study the variation in topology in protein structure space. The consistent observation of 80/20 rule in topology space is perceptible as shown by Figure 1 and Figure S5. This can be drawn in parallel to “Pareto-distribution” that is eminent in across fields of natural sciences and economics [21, 22]. The variance in protein space in “Prevalent” and “Non-prevalent” groups are majorly influence by the nature of “structure class”. However, the emergent pattern of “small fraction of topology mostly populating structure space” can be drawn.

### Topology visualization

ProLego draws protein topology diagrams using “Topology String”. The pipeline extracts nodes (SSEs) and edges (sequential and spatial contact) using developed topology visualizer and renders in 2D and 1D SVG plots (Supplementary Section 1.6 and 1.7). Figure 1A, shows the ProLego result for photosynthetic reaction centre protein (PDB id: 1JB0, chain L). The protein chain has N-terminal β-hairpin, followed by seven α-helices. As shown by the Figure 1A, ProLego generates (i) secondary structure contact map, (ii) 2D-cartoon view and (iii) linear topology graphs, representing different ways to examine protein topology. The secondary structure contact map illustrates the presence of contact and their relative orientation with different colour codes. Similar colour codes have been used in the linear topology which shows secondary structures from N to C terminal with strands as triangles (up/down relative to orientation) and rectangle blocks as alpha helices. Spatial contacts between SS have been shown as arcs. The cartoon view, illustrates the protein topology graph where solid lines show the sequential SSE contact whereas, the dashed red line shows the presence of tertiary contact between corresponding secondary structures.

### Extracting protein topology modules

This protocol extracts sub-structures or modules from a protein by analysing topology string. Fixing a window of one SSE from N to C-terminal, all observable protein topology in a chain has been listed (Figure S1, Supplementary Section 1.3). For a protein with “n” SSE (n>3), the search extracts “n-1” SSE-topology modules, following the SSE combination stepwise from N to C terminal. The topology database, ProLegoDB, are then used to map the protein chains and domains with each resultant topology modules. An example of extracted topological modules has been listed in Table S3.

### Topology Database

ProLegoDB is an extensive database for protein topology. The database is the collection of unique topologies extracted from non-redundant protein sets, generated from PDB (using PISCES-server [23]) and curated domain databases (Supplementary Section 1.5). This database has 58,186 protein chains and 14408 protein domains topology analysed and grouped into 7201 statistically significant topology groups (Table 1). As the topologies are defined as per their secondary structure construct, its relatively easy to divide the whole space into all-alpha (A), all-beta (B) and alpha-beta (AB), structure classes. Each topology has been reported with observed occurrence frequency and statistical significance score (Supplementary section 15)

A search in ProLegoDB can be performed from three levels i.e. Topology, Protein and Domains. Using “Search by Topology”, user can provide queries as per SS composition or advanced query of filtering with numbers of helix, strands as well as statistical significance. The query result lists all possible topologies with requested SS-composition along with their significant scores. Each row of the result has the corresponding link describing topology.

### Comparison of ProLego with PTGL

Among current state-of-the-art protein graph generation servers (Table S1), PTGL is the most recent [17]. This is a subsequent upgrade and development over protein topology graph library [24, 25]. PTGL’s integration of graph modelling language (GML) for visualization is one of the first kind to apply in protein graphs (Figure 1B). The most recent addition include ligand information in protein secondary structure contact and decomposes protein chain into alpha, beta and alpha-beta and receptor–ligand graphs[17]. The approach is shown to be used for searching sub-graphs, which is a crucial aspect of protein graph analysis, as also reported by Pro-Origami [16] and Tableaux [26].

Both PTGL and ProLego, address the topological graph from secondary structures. A comparative study on type of topology visualisation for PTGL and ProLego has been shown in Figure 1. With ProLego, we illustrate the usability of string based topological representation. ProLego, provides more detailed and modular view to protein topology landscape. Our primary focus is to describe the variation in protein topology space, hence have not considered the ligand interactions. However, in the context of protein topology, ProLego provides topological frequencies (as P/Np) and statistical significance for all reported topologies. The extensive topology database, with different search modules, is advantageous to tailor search for topology. Identification of topological modules remains one of the most significant development in ProLego as compare to other topology databases. Use of topology string, not only increases the computational efficiency but also provides more intuitive result (supplementary section 1.8).

## Conclusion

With ProLego, we aim to provide an alternative approach to study protein structure topology. ProLego is inspired by modular architecture in protein topology space, which can be easily studied by the proposed “Topology String”. The component approach is found to be efficiently scanning the structure space and explore the nature of topology space. To understand the secondary structure based architecture in proteins, ProLego have compiled an extensive topology database analyzing different sets of non-redundant representative protein datasets. The server application provides an easy access to the database as well as enables users to investigate their protein of interest. With the integration of state-of-the-art framework and libraries, improved topology visualization approaches have been implemented and compared with other open source topology servers. Exclusively, ProLego-Server can be used for identifying constituent topological modules in proteins of interest, which could be used as “lego-blocks” in protein designing.

## Availability and requirements

Project home page: <u>http://www.proteinlego.com/</u>

Operating system(s): Platform independent Programming language: Python, JavaScript

Other requirements: Morden web-browser (Chrome, Firefox, Safari updated after June 2016) License: FreeBSD etc.

Any restrictions to use by non-academics: none

**List of abbreviations**
SSE: Secondary Structure Elements
P: Prevalent topology
NP: Non-Prevalent topology
PDB: Protein Data Bank

## Declarations

Ethics approval and consent to participate: Not applicable Consent for publication: Not applicable Availability of data and martial: Studied protein datasets are listed in the project site <u>http://www.proteinlego.com</u>

Competing interests: None

Funding: TK has been supported by UGC-MANF SRF, DBT-CCPM, DBT-CoE. SKP has been supported by DBT-BINC JRF. The funding bodies have no role in study, designing or conducting the experiments.

Authors’ contributions: TK and SKP equally contributed to designing and building the application. IG and TK have developed the algorithm. TK, SKP and IG wrote the manuscript. Acknowledgements: Authors will like to thanks all group member of Prof. Ghosh and to Dr. Rama Kaalia for valuable input and improving the manuscript.

